# Updating genome annotation for the microbial cell factory *Aspergillus niger* using gene co-expression networks

**DOI:** 10.1101/398842

**Authors:** p Schäp, MJ Kwon, B Baumann, B Gutschmann, S Jung, S Lenz, B Nitsche, N Paege, T Cairns, V Meyer

**Author notes:** equally contributed to this work (p Schäp, MJ Kwon).

## Abstract

A significant challenge in our understanding of biological systems is the high number of genes with unknown function in many genomes. The fungal genus Aspergillus contains important pathogens of humans, model organisms, and microbial cell factories. *Aspergillus niger* is used to produce organic acids, proteins, and is a promising source of new bioactive secondary metabolites. Out of the 14,165 open reading frames predicted in the *A. niger* genome of only 2% have been experimentally verified and over 6,000 are hypothetical. Here we show that gene co-expression network analysis can be used to overcome this limitation. A meta-analysis of 155 transcriptomics experiments generated co-expression networks for 9,579 genes (∼65%) of the *A. niger* genome. By populating this dataset with over 1,200 gene functional experiments from the genus *Aspergillus* and performing gene ontology enrichment, we could infer biological processes for 9,263 of *A. niger* genes, including 2,970 hypothetical genes. Experimental validation of selected co-expression sub-networks uncovered four transcription factors involved in secondary metabolite synthesis, which were used to activate production of multiple natural products. This study constitutes a significant step towards systems-level understanding of *A. niger*, and the datasets can be used to fuel discoveries of model systems, fungal pathogens, and biotechnology.

## Introduction

The genus Aspergillus (phylum Ascomycota) is comprised of nearly 200 species of saprophytic and ubiquitous fungi, and includes important pathogens of humans (*Aspergillus fumigatus*), model organisms (*Aspergillus nidulans*) and microbial cell factories (*Aspergillus oryzae, Aspergillus niger*). *A. niger* has been exploited for over a century by biotechnologists for the production of organic acids, proteins and enzymes (1). It is the major worldwide producer of citric acid with an estimated value of $2.6 billion in 2014, which is predicted to rise to $3.6 billion by 2020 (2). As a prolific secretor of proteins, *A. niger* is used to produce various enzymes at a bulk scale (1). The first *A. niger* genome was sequenced in 2007, which contained an estimated 14,165 coding genes about 6,000 of which were hypothetical (3). More recent sequencing of additional *A. niger* strains and other genomes from the genus *Aspergillus* (4,5), combined with refinement of online genome analyses portals (6–8), and comparative genomic studies amongst the Aspergilli (5), have not significantly increased the percentage of *A. niger* genes that have functional predictions. While the exact number of ‘hypothetical’ genes varies between databases and *A. niger* genomes, recent estimates suggest that between 40 – 50% of the genes still remain hypothetical (1,9). Furthermore, only 2% of its genes (247) have a verified function in the Aspergillus Genome Database (AspGD (10)). Even for the gold-standard model organism *Saccharomyces cerevisiae,* 21% of its predicted genes remain dubious (11), despite its high genetic tractability and a research community with more than 1,800 research labs worldwide. Indeed, gene functional predictions for *A. nidulans, A. fumigatus, A*. *oryze* and other Aspergilli typically cover 40-50% of the genome (9,10)

Such high frequency of unknown and hypothetical genes severely limits the power of systems-level analyses. One approach to overcome this limitation involves the generation and interrogation of gene expression networks based on transcriptomic datasets (12–14). The hypothesis underlying this approach is that genes which are robustly co-expressed under diverse conditions are likely to function in the same or in closely related biological processes (15). In this study, we thus conducted a meta-analysis of 155 publically available transcriptomics analyses for *A. niger*, and used these data to generate a genome-level co-expression network and sub-networks for more than 9,500 genes. To aid user interpretations of gene biological process, gene sub-networks were analysed for enriched gene ontology (GO) terms, and integrated with information gleaned from 1,200 validated genes from the genus *Aspergillus*. Interrogation of selected co-expression sub-networks for verified genes and randomly selected hypothetical genes confirmed high quality datasets that enable rapid and facile predictions of biological processes. This co-expression resource will be integrated in the functional genomic database FungiDB (6) for use by the research community.

We finally validated this co-expression network by selecting two co-expression sub-networks predicted to be involved in natural product synthesis of *A. niger*. The rationale behind was that (i) there is an urgent need for new drugs due to the emergence of multiresistant bacteria and fungi (16), (ii) most of the natural product repertoire of filamentous fungi such as *A. niger* is unknown(17) and (iii) *A. niger* has been shown to be a superior expression host for medicinal drugs in g/L scale (18). Experimental validation included generation of null and overexpression mutants of transcription factors present in these sub-networks, controlled bioreactor cultivations and global analysis of gene expression at transcript and metabolite level. The co-expression resources and experimental validation developed in this study thus enable high quality gene functional predictions in *A. niger*.

## Methods

### Strains and molecular techniques

Media compositions, transformation of *A. niger*, strain purification and fungal chromosomal DNA isolation were described earlier (19). Standard PCR and cloning procedures were used for the generation of all constructs (20) and all cloned fragments were confirmed by DNA sequencing. Correct integrations of constructs in *A. niger* were verified by Southern analysis (20). In the case of overexpressing TF1, TF2 and HD, the respective open reading frames were cloned into the Tet-on vector pVG2.2 (21) and the resulting plasmids integrated as single or multiple copies at the *pyrG* locus. Deletion constructs were made by PCR amplification of the 5’- and 3’-flanks of the respective open reading frames (at least 0.9 kb long). N402 genomic DNA served as template DNA. The histidine selection marker (22) was used for selecting single deletion strains, whereas the *pyrG* marker was used for the establishment of the strain deleted in both TF1 and TF2. Details on cloning protocols, primers used and Southern blot results can be requested from the authors.

### Gene network analysis and quality control

As of March 02, 2016, 283 microarray data (platform: <u>GPL6758</u>) of *A. niger* covering 155 different cultivation conditions were publically available at the GEO database(23), whose processing and normalization of the arrays have been published (24). In brief, array data in the form of CEL-files (25) were processed using the Affymetrix analysis package (25) (version 1.42.1) from Bioconductor (26) and expression data were calculated for genes under each condition with an MAS5 background correction. Pairwise correlations of gene expression between all *A. niger* genes were generated by calculating the Spearman’s rank correlation coefficient (27) using R. To assess a cut-off indicating biological relevance, Spearman correlations were firstly calculated using a pseudo random data set whereby normalized transcript values for each individual gene were randomized amongst the 283 arrays and 155 experimental conditions. Using this pseudo random data set, the Spearman rank coefficient was calculated pairwise for all predicted *A. niger* genes, giving a total of 104,958,315 comparisons/calculated spearman rank coefficients, from which 52,476,536 were positively correlated. From these, only 2 were greater than |0.4|and none above |0.5|. Subsequently |0.5| was taken as a threshold for co-expression. Sub-networks were calculated at an individual gene level using R. All genes that were co-expressed with individual query ORFs will be reported at FungiDB (6) and using a |≥0.5| and |≥0.7| Spearman cut-off.

To expedite investigation of the sub-networks and their common biological process, gene ontology enrichment (GOE) was implemented using Python version 2.7.13. GO terms and their hierarchical structure were downloaded from AspGD(10). Enriched GO Biological Process terms for all genes residing in a query sub-network were calculated relative to the *A. niger* genome and statistical significance was defined using the Fishers exact test (p-value <0.05). For informant ORFs, experimentally verified *A. niger* genes were retrieved from AspGD (10). Additionally, *A. niger* orthologs for any gene with wet lab verification in *A. fumigatus, A. nidulans* or *A. oryzae* were identified using the ENSEMBL BLAST tool using default settings (8). Finally, 81 secondary metabolite core enzymes (28,29) were also defined as informant ORFs. We generated informant ORF (‘prioritized ORF’) sub-networks, which report significant co-expression of query genes exclusively with one or more informant ORFs.

### Reporter gene expression

Protocols for luciferase-based measurement of gene expression in microtiter format based on Tet-on (21) or *anafp* (24) promoter systems have been published. In case of strain BBA17.6, unable to form spores, the strain was inoculated on complete medium and allowed to grow for 7 days at 30°C. Biomass was harvested using physiological salt solution and used for inoculation. All data shown derived from biological duplicates each measured in technical quadruplicates if not otherwise indicated. Raw datasets can be requested from the authors.

### Bioreactor cultivation

Medium composition and the protocol for glucose-limited batch cultivation of *A. niger* in 5L bioreactors have been described (21). In the case of strains overexpressing TF1 (strain MJK10.22) and TF2 (strain MJK11.17), the Tet-on system was induced with a final concentration of 10 µg/ml doxycycline when the culture reached 1 g/kg dry biomass. Samples for transcriptional and metabolome profiling were taken ∼6 h (∼72 h) after induction, i.e. during exponential (post-exponential) growth phase. To ensure constant activation of the Tet-on system throughout cultivation, 10 µg/ml doxycycline was added every 10-12 hours (five times in total). For control (MJK17.25) and deletion strains (MJK14.7, MJK16.5, MJK18.1), doxycycline was added twice; once after the culture reached 1 g/kg dry biomass and 24 hours before samples were taken from post-exponential growth phase for transcriptomics and metabolomics analyses.

### Transcriptional profiling

Total RNA extraction, RNA quality control, and RNA sequencing were performed at GenomeScan (Leiden, the Netherlands). Quality analysis of raw data was done as previously described (30). In brief, ∼13 million reads of 150 bp were obtained from paired-end mode for each sample. Read data were trimmed and quality controlled with FastQC (http://www.bioinformatics.babraham.ac.uk/projects/fastqc/). STAR (31) was used to map the reads to the *A. niger* CBS 513.88 genome (http://fungi.ensembl.org/). On average, the unique alignment rate was ∼95%. Data normalization was performed with DEseq2 (32). Differential gene expression was evaluated with Wald test with a threshold of the Benjamini and Hochberg False Discovery Rate (FDR) of 0.05 (33) with DEseq2.

## Metabolome profiling

Metabolites were extracted from biomass corresponding to 2.5 mg biomass dry weight by Metabolomic Discoveries GmbH (Potsdam, Germany) and identified based on Metabolomic Discoveries’ database entries of authentic standards. Liquid chromatography (LC) separation was performed using hydrophilic interaction chromatography with a iHILIC®-Fusion, 150x2.1 mm, 5 µm, 200 Å 5 μm, 200 A column (HILICON, Umeå Sweden), operated by an Agilent 1290 UPLC system (Agilent, Santa Clara, USA). The LC mobile phase was (i) 10 mM ammonium acetate (Sigma-Aldrich, USA) in water (Thermo, USA) with 5% and 95% acetonitrile (Thermo, USA) (pH 6) and (ii) acetonitrile with 5% 10 mM ammonium acetate in 95% water. The LC mobile phase was a linear gradient from 95% to 65% acetonitrile over 8.5 min, followed by linear gradient from 65% to 5% acetonitrile over 1 min, 2.5 min wash with 5% and 3 min re-equilibration with 95% acetonitrile. The flow rate was 400 μl/min and injection volume was 1 μl. Mass spectrometry was performed using a high-resolution 6540 QTOF/MS Detector (Agilent, Santa Clara, USA) with a mass accuracy of < 2 ppm. Spectra were recorded in a mass range from 50 *m/z* to 1700 *m/z* at 2 GHz in extended dynamic range in both positive and negative ionization mode. The measured metabolite concentrations were normalized to the internal standard. Significant concentration changes of metabolites in different samples were analyzed by appropriate statistical test procedures (ANOVA, paired t-test) using R. When the adjusted p value based on Benjamini and Hochberg FDR(33) was lower than 0.05 and the fold change (log2) higher than +/-1, expression of the metabolites was considered as significantly different.

## Results

### The transcriptomic landscape of *A. niger* inferred from a gene expression meta-analysis

We normalized and interrogated gene expression across 155 published transcriptomic analyses for the *A. niger* laboratory wildtype strain N402 (ATCC 64974) and its descendants, comprising of 283 Affymetrix microarray experiments in total (24). Experimental parameters include a diverse range of cultivation conditions (agar plate, bioreactor, shake flask), developmental and morphological stages (germination, mycelial growth, sporulation), deletion and disruption mutants, stress conditions (antifungals, secretion stress, pH), different carbon and nitrogen sources, starvation, and co-cultivation with bacteria. These experimental conditions represent diverse niches inhabited by *A. niger* as well as industrial cultivation conditions, in addition to (a)biotic and genetic perturbations that result in global changes in gene expression.

In order to demonstrate that accurate values of transcript abundance were derived from this meta-analysis, we plotted average gene expression values for each gene throughout the 155 conditions as a function of chromosomal locus (Fig. 1A). From these data, we categorized low, medium, and highly expressed loci. Subsequently, we generated a DNA cassette expressing a luciferase reporter gene under control of the inducible Tet-on promoter (21) and targeted it to the 5-upstream region of two low and one high expression locus (Fig. 1B). The *pyrG* locus present on chromosome III, routinely used for gene-targeted integration in *A. niger*, served as locus control for Tet-on driven medium expression of luciferase (21). Luciferase levels measured at these loci were confirmed to be low, medium and high in relative terms in microtiter cultivations of the different *A. niger* strains (Fig. 1C) and in controlled batch cultivation at bioreactor scale (Fig. 1D). We therefore conclude that the microarray data accurately reflect *A. niger* gene expression values. Note that this transcriptomic landscape is a significant addition to the *A. niger* molecular toolkit, as it facilitates rational control of gene dosage (time of induction and absolute expression level) by targeted locus-specific integration of a gene of interest.

**Figure 1.**
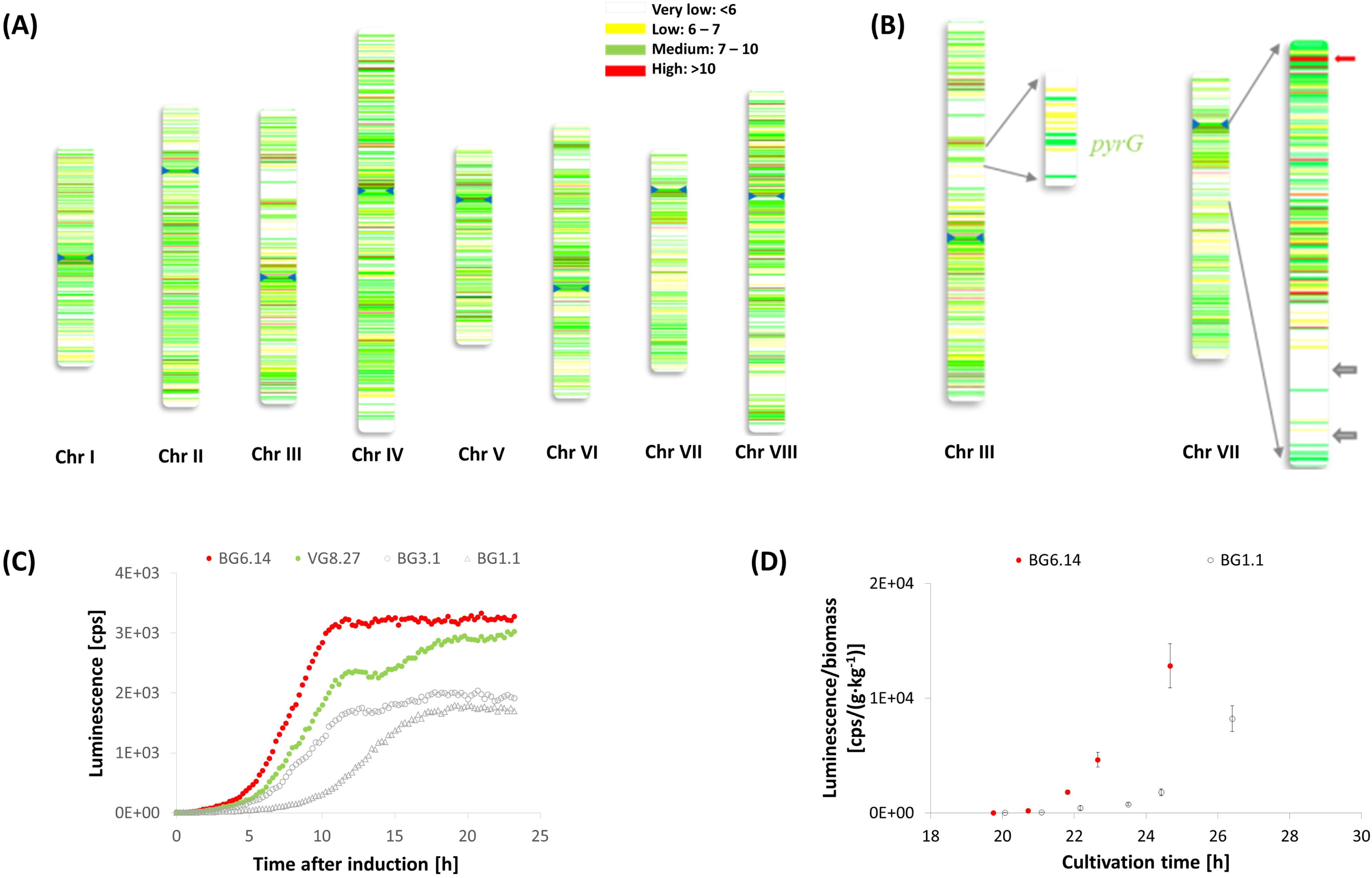
The transcriptomic landscape of *A. niger*. (A) Chromosomal expression values expressed as log2 mean from 155 cultivation conditions. Blue arrows indicated centromere position on each chromosome(3). (B) Selected chromosomal positions in chromosome III and VII for *in vivo* verification. Red arrow indicates position of chosen integration site at high level expression locus, white arrows indicate positions of chosen integration sites at low level expression loci, respectively. (C) Data for luciferase activity measured by luminescence in counts per second (cps) over time after induction in hours in microtiter plates. Data for strains expressing luciferase at the locus An11g08480 (‘low gene expression locus’; strain BG1.1), An09g02630 (‘low gene expression locus’; strain BG3.1), An12g03570 (‘medium gene expression locus’, *pyrG*, strain VG8.27), and An11g11310 (‘high gene expression locus’; strain BG6.14) are given. Doxycycline was added 16 h after inoculation. (D) Cultivation of strain BG1.1 and BG6.14 in 5L bioreactors. For each strain, two batch cultivations were performed; one was induced with 5 µg/ml doxycycline at a dry biomass concentration of 2 g/kg. Cultivation in the presence of doxycycline mediated gene expression as demonstrated by increased luminescence after induction measured in technical quintuplicate.

### Construction of high quality co-expression networks for *A. niger*

Experimentally validated gene expression data from the *A. niger* transcriptional meta-analysis was utilized to generate a gene co-expression network based on Spearman’s rank correlation coefficient (27). In order to define a minimum Spearman’s rank correlation coefficient (ρ) for which we could be confident in extracting biologically meaningful co-expression, we conducted a preliminary quality control experiment, where transcript values for each individual gene were randomized amongst the 155 experimental conditions. This gave a dataset with identically distributed but randomized expression patterns. Next, we calculated every possible transcriptional correlation between genes on the *A. niger* genome, resulting in over 100 million ρ-values. This identified 52 million positive and 48 million negative correlations (Fig. 2). From this dataset, only 2 ρ-values were above |0.4|, and none were above |0.5|. Consequently, we took ρ ≥ |0.5| as a minimum cut-off for biologically meaningful co-expression relationships. Calculations of Spearman correlations using the non-randomized microarray data resulted in over 4.5 million correlations which passed the minimum ρ ≥ |0.5| cut-off. From these datasets, co-expression sub-networks for every gene in the global network were generated for both positively and negatively correlated genes (Fig. 2). We classified them into two groups: ‘stringent’ (ρ ≥ |0.5|, 9,579 genes) and ‘highly stringent’ (ρ ≥ |0.7|, 6,305 genes) and calculated enriched GO terms for each gene sub-network relative to the *A. niger* genome.

**Figure 2:**
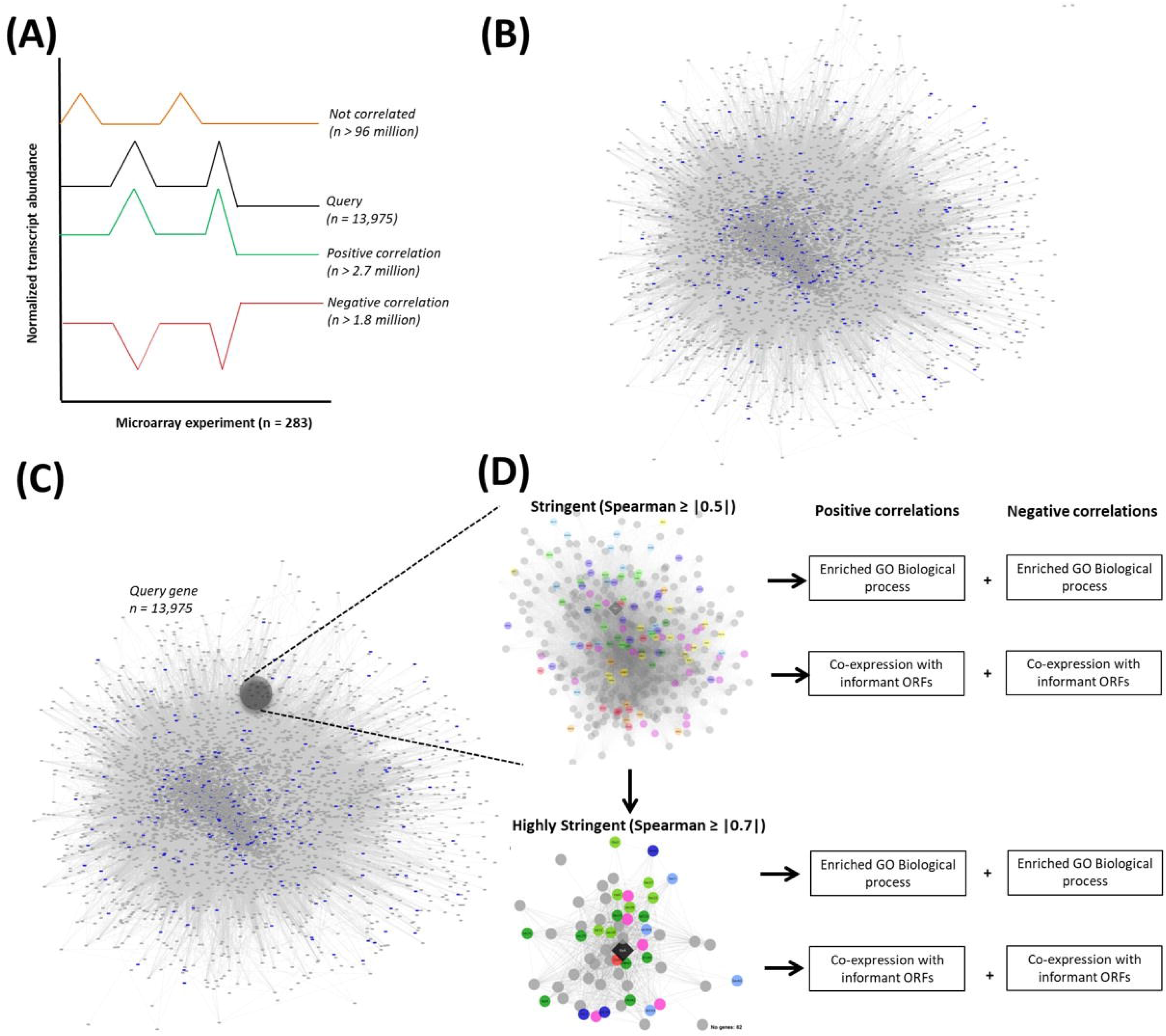
Workflow to generate gene co-expression resources. (A) Various transcriptional signatures across 155 cultivation conditions of *A. niger* obtained from 283 microarrays are schematically represented. Spearman’s rank correlation coefficients were calculated pairwise between all remaining predicted genes in the *A. niger* genome. Over 96 million correlations between gene pairs were not defined as co-expressed based on the |0.5|cut-off (black and orange lines), whereas 2.7 million were positively correlated (black and green lines) and 1.8 million were negatively correlated (black and red lines). These >4.5 million significant co-expression relationships are visualized as a network in (B), with genes shown as grey squares, and correlations as lines. The length of each line is proportional to the Spearman correlation between gene pairs. Genes predicted to encode transcription factors(3) are highlighted in blue. (C) For each individual gene in the network (13,975 in total; this number deviates from 14,165 reported in 2007(3), because we have omitted truncated ORFs reported there from our analysis), sub-networks were calculated which report all significant correlations with the query gene of ρ ≥ |0.5| (stringent dataset) and of ρ ≥ |0.7| (very stringent dataset). Both sub-networks were analyzed for significantly enriched GO Terms (biological processes). Additionally, we highlight co-expression between query genes and ‘informant ORFs’, which have either been experimentally validated in the Aspergilli, or are predicted to encode key secondary metabolite biosynthetic enzymes (D).

### Integration of co-expression networks with community-wide experimental evidence of gene function

The *Aspergillus* community has functionally characterized over a thousand of genes in different species of the genus *Aspergillus*, which we reasoned can be used to aid *a priori* predictions of hypothetical genes or not yet verified genes in *A. niger*. In order to integrate such experimental data with the co-expression network, we mined the *Aspergillus* genome database AspGD (10) to generate a near-complete list of ORFs that have been functionally characterized in Aspergilli. All experimentally validated ORFs for *A. niger* (n = 247), *A. nidulans* (639), *A. fumigatus* (218) and *A. oryzae* (81) were included in this dataset. Given the strong potential of *A. niger* as a platform for discovery and production of new bioactive molecules, we also included 81 putative polyketide synthase (PKS) or nonribosomal peptide synthetase (NRPS) encoding genes of *A. niger* that reside in 78 predicted secondary metabolite clusters (28,29), giving in total 1,266 prioritized ORFs. For every gene in the *A. niger* genome, we calculated co-expression interactions specifically with these 1,266 rationally prioritized ORFs. A total of 9,263 (ρ ≥ |0.5|) and 5,178 (ρ ≥ |0.7|) candidate genes had one or more correlations with prioritized ORFs. These datasets thus constitute the most comprehensive co-expression resource for a filamentous fungus and will be accessible at FungiDB(6).

### Co-expression resources enable facile predictions of gene biological function

In order to test whether biologically meaningful interpretations of gene and network function can be extracted from these resources, we interrogated both stringent and highly stringent datasets for genes where the biological processes, molecular function, and subcellular localization of encoded proteins have been studied in fungi and which represent the broad range of utilities and challenges posed by fungi. From the perspective of industrial biotechnology, we interrogated networks for the gene encoding the ATPase BipA, which is required for high secretion yield of industrially useful enzymes by acting as chaperone to mediate protein folding in the endoplasmic reticulum (34). With regards to potential drug target discovery, we analyzed gene expression networks for Erg11 (Cyp51), which is the molecular target for azoles (35). For assessment of virulence in both plant and human infecting fungi, we interrogated networks for the NRPS SidD, which is necessary for the biosynthesis of the siderophore triacetyl fusarinine C, and ultimately iron acquisition during infection (36). Assessment of all control sub-networks at GO and individual gene-level revealed striking co-expression of genes encoding proteins involved in respective metabolic pathways, associated biological processes, subcellular organelles, protein complexes, known regulatory transcription factors/GTPases/chaperones, and cognate transporters, amongst others (Fig. 3). The lowest Spearman correlation coefficient of |0.5| clearly results in biologically meaningful gene co-expression as exemplified by the delineation of diverse yet related processes, including orchestration of retrograde/anterograde vesicle trafficking via COPI/COPII/secretion associated proteins (BipA) (37), coordination of ergosterol biosynthesis by sterol regulatory binding element regulators SrbA/SrbB and association of this pathway with respiration at the mitochondrial membrane (Erg11) (38), and the linking of respective ergosterol and ornithine primary and secondary metabolic pathways during siderophore biosynthesis via the interdependent metabolite mevalonate (SidD) (39). With regards co-expressed genes as a function of chromosomal location, a hallmark of filamentous fungal genomes is that genes necessary for the biosynthesis of secondary metabolite products occur in physically linked contiguous clusters. SidD resides in a six-gene cluster with SidJ, SidF, SidH, SitT and MirD, all of which were represented in the high stringency network that contained a total of only 13 genes (Fig. 3).

**Figure 3:**
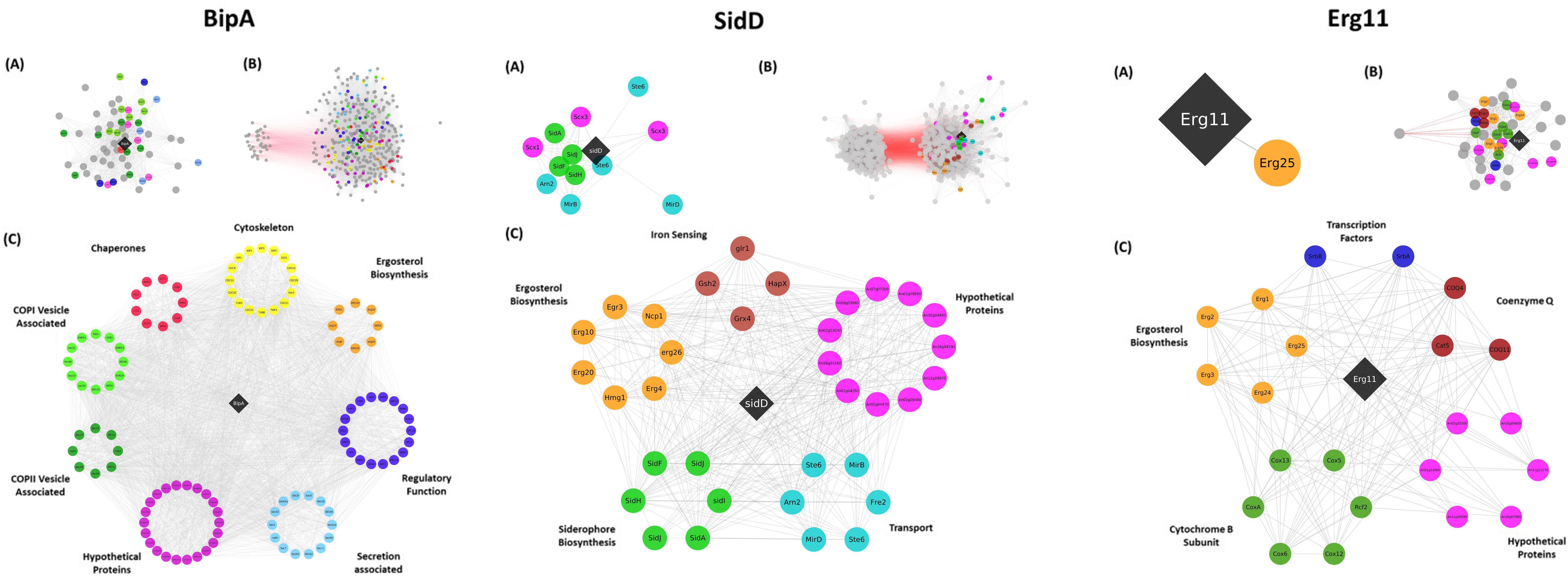
Schematic representation of co-expression networks for BipA, SidD, and Erg11 encoding genes. Genes are represented by circles, with positive and negative correlations depicted by grey and red lines, respectively. For simplicity, protein names are given in the centre of each circle. Where names were not available in *A. niger*, we used the name for either *A. nidulans* or *S. cerevisiae* ortholog. Query genes encoding BipA, SidD, or Erg11 are given in black diamond boxes. Co-expression sub-networks are given for gene pairs passing |0.7| (A) and |0.5| (B) Spearman correlation coefficient cut-offs. Both sub-networks were assessed for enriched GO terms, and genes were sorted into functional categories based on these analyses and manual interrogation of research literature (C). Note that in each instance, hypothetical proteins were co-expressed with each query gene, indicating the encoded products are somehow associated with these biological processes.

Based on enriched GO terms from gene sub-networks and co-expression with experimentally verified ORFs, we could further rapidly infer biological processes for a total of 2,970 (ρ ≥ |0.5|) and 1,016 (ρ ≥ |0.7|) hypothetical genes that were positively and/or negatively associated using this analysis. Additionally, we interrogated entire families of functionally related genes that have been well characterized in the Aspergilli, including phosphatases, chromatin remodelers, and transcription factors and were able to assign novel biological processes for all these predicted genes. Taken together, these quality control experiments strongly suggest that the co-expression resources developed in this study can be used for high confidence hypothesis generation at a variety of conceptual levels, including biological process, metabolic pathway, protein complexes, and individual genes.

### Co-expression resources accurately predict transcription factors of the ribosomally synthesized natural product AnAFP of *A. niger*

In order to provide experimental confirmation in predictions of biological processes gleaned from this co-expression resource, we interrogated all datasets associated with the gene encoding the *A. niger* antifungal peptide AnAFP. Ribosomally synthesized antifungal peptides of the AFP family are promising molecules for use in medical or agricultural applications to combat human- and plant-pathogenic fungi(40). We and others could show that expression of their cognate genes are under tight temporal and spatial regulation in their native hosts and precedes asexual sporulation (24,41–43).

The gene encoding AnAFP (An07g01320), is co-expressed with 986 genes (ρ ≥ |0.5|; 605 positively correlated / 381 negatively correlated (24)). GO enrichment analyses of positively correlated sub-networks uncovered that *anafp* gene expression parallels with fungal secondary metabolism, carbon limitation and autophagy (24). In total, 23 predicted transcription factors are co-expressed at a stringent level among which were the transcription factors VelC (An04g07320) and StuA (An05g00480; Fig. 4), both of which are key regulators of asexual development and secondary metabolism in Aspergilli (44–46). In order to confirm a regulatory function of these transcription factors on *anafp* expression, we used a reporter strain in which the *anafp* ORF has been replaced with a luciferase gene. Deletion of *stuA* or *velC* in this background revealed a strong increase or decreased/delayed activation of the *anafp* promoter, respectively (Fig. 4). Interestingly, the transcription factor binding site for VelC is unknown, whereas the binding sequence for StuA is absent from the predicted promoter region of *anafp*. These data thus indicate that the resources generated in this study enable accurate predictions of (in)direct regulatory proteins even in the absence of DNA binding sites.

**Figure 4:**
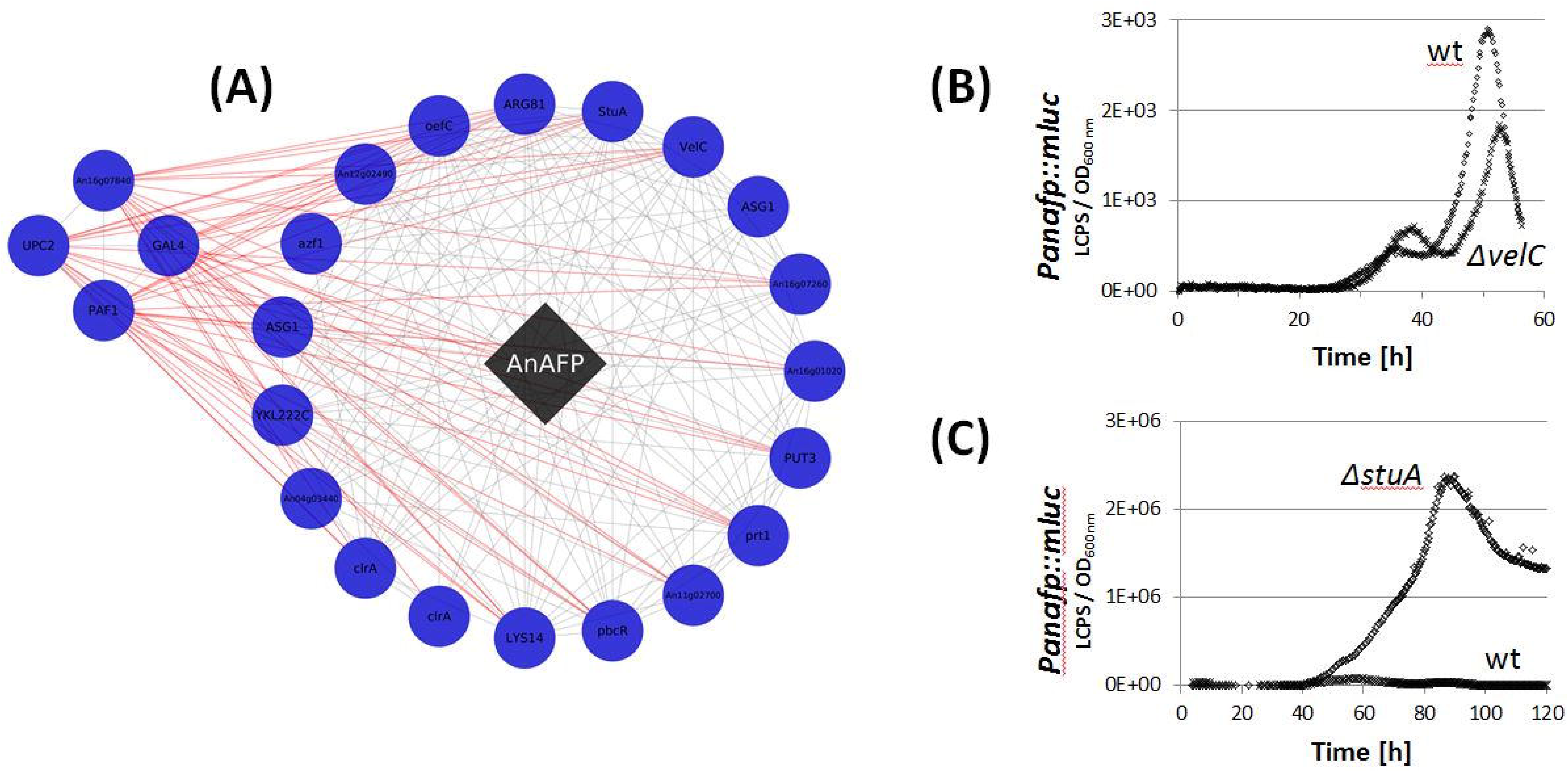
*In vivo* proof of the predictive power of gene-coexpression sub-networks. (A) Depicted are 23 predicted transcription factors in the *anafp* sub-network which either positively (19) or negatively (4) correlate with the expression of the *anafp* gene. Two of these (*velC* and *stuA*) were selected for *in vivo* verification by establishing *velC* and *stuA* deletion strains in an *anafp::mluc* reporter strain published earlier (strain PK2.9)(24). (B, C) Analysis of *anafp* promoter activity in a strain deleted for *velC* (strain BBA17.9) and for *stuA* (strain BBA13.2), respectively, which was compared to the progenitor strain PK2.9 abbreviated as ‘wt’. Data are from liquid microtiter plate cultures, where a defined amount of biomass was used for inoculation and luciferase expression monitored online during cultivation. Expression levels are depicted for one representative example from three independent experiments. Each experiment was performed in quintuplicate.

### Co-expression resources accurately predict transcription factors of non-ribosomally synthesized natural products of *A. niger*

The transcriptional activation of secondary metabolite (SM) gene clusters in different filamentous fungi is one current focus of the fungal research community (9) as more than 60% of currently approved clinical drugs are derived from natural products (47). *A. niger* stands out due to its exceptional high number of predicted SM gene clusters in its genome (78), harboring 81 core enzymes in total, such as NRPS and PKS (28). However, only a dozen of SMs have been identified from *A. niger* so far (48). Our survey of the expression data of all gene clusters under the 155 cultivation conditions uncovered that the majority of SM core genes (53) are expressed in at least one condition. The majority of expressed core genes are also co-expressed with their cluster members. Notably, not all cluster members are co-expressed with contiguous transcription factors. Indeed, only ∼30% of the gene clusters display co-expression with contiguous transcription factors. We thus questioned which transcription factors are regulating these SM gene clusters, and used the co-expression dataset to assign biological processes to genes predicted to encode transcription factors. Given the important role of chromatin remodelers in activation and silencing of secondary metabolite clusters, we also interrogated genes predicted to encode histone deacetylases (49). This identified two ORFs encoding putative transcription factors: An07g07370 (TF1) and An12g07690 (TF2), and a histone deacetylase (An09g06520, HD) that are positively and negatively (TF1, TF2) or only negatively (HD) co-expressed with numerous core SM genes. Notably, all three genes do not reside in contiguous SM gene clusters but belong to a large SM sub-network consisting of 152 genes including 26 SM core genes, whereby gene expression of TF1 and TF2 correlate very strongly (ρ = 0.87). Interrogation of enriched GO terms for both TF1 and TF2 gene sub-networks revealed enrichment of fatty acid metabolism, autophagy, mitochondria degradation (positively correlated) and maturation of rRNA and tRNA, ribosomal assembly and amino acid metabolism (negatively correlated). This analysis thus allowed us to select genes for *in vivo* functional studies based on a non-intuitive selection procedure. Neither TF1, TF2 nor HD have been experimentally characterized in fungi so far. In order to confirm a regulatory function of these putative regulators on SM core gene expression, we generated (i) single deletion strains for TF1, TF2 and HD, respectively, (ii) a double deletion strain for TF1 and TF2, and (iii) individual conditional overexpression mutants for TF1, TF2 and HD using the Tet-on system (21). The strongest effect on the metabolome profile of *A. niger* was observed during overexpression of TF1 and TF2 (Fig. 5). SMs up-regulated under these conditions included, but were not limited to aurasperones, citreoviridin D, terrein, aspernigrin A, nigerazine A and B, pyranonigrin A and D, flavasperone, fonsecin, O-demethylfonsecin, flaviolin, funalenone; among the SMs down-regulated under these conditions were asperpyrone, L-agaridoxin and nummularine F (Fig. 5). These physiological changes were paralleled by up/down-regulation of thousands of genes as determined by RNA-Seq analyses, whereby controlled overexpression of TF1 (TF2) modulated expression of 45 (43) SM core genes especially during post-exponential growth phase of *A. niger* (Fig. 5). This strongly suggests that both transcription factors are likely global regulators modulating gene expression dynamics during late growth stages of *A. niger* either directly or indirectly.

**Figure 5:**
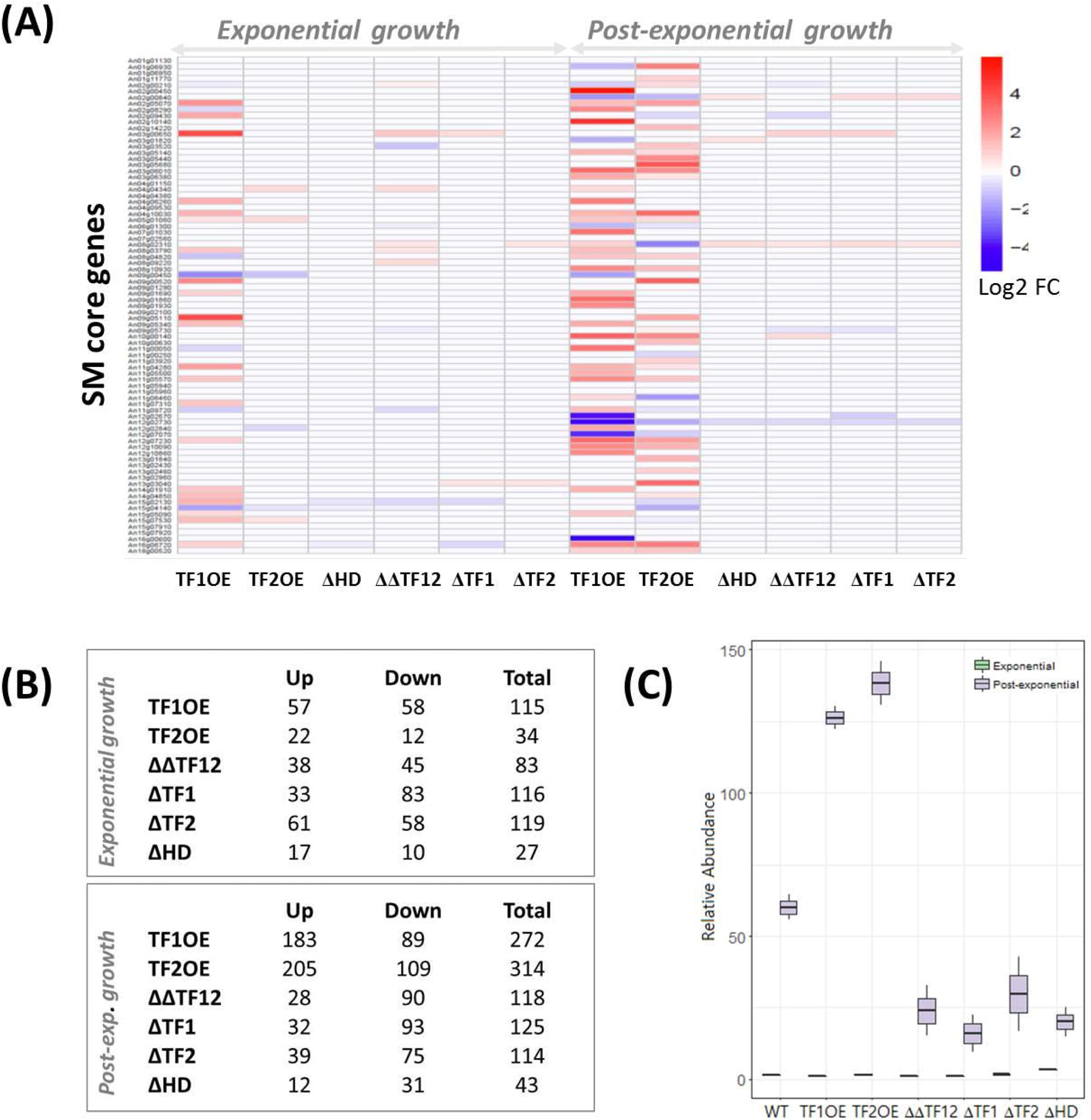
Co-expression sub-networks uncover global regulators of secondary metabolism in *A. niger*. Controlled batch cultivations using 5L bioreactor systems were performed with the progenitor strain and strains deleted for HD, TF1 and/or TF2 (ΔTF1, ΔTF2, ΔΔTF12, ΔHD) or overexpressing either transcription factors (OETF1, OETF2). Note that we excluded the strain overexpressing HD from this analysis because its gene expression is negatively correlated with the mostly silent SM core genes. Samples were taken for untargeted metabolome analyses (LC-QTOF/MS) and global RNA-Seq during exponential and post-exponential growth phase. Note that the maximum growth rate was for all strains 0.24 h^-1^ except for TF1OE (0.19 h^-1^). (A) Differential gene expression for 81 predicted secondary metabolite core genes in mutant isolates relative to the progenitor control as determined by RNA-Seq analysis. (B) Assessment of metabolite abundance demonstrated differential abundance of hundreds of *A. niger* metabolites following deletion or over-expression of HD, TF1 and TF2. In total, 3,410 primary and secondary metabolites were detected in all runs, 1,260 of which were known compounds and 478 thereof were differentially expressed upon deletion or overexpression of these putative regulators. (C) Relative abundance of the SM aurasperone B is shown in progenitor control and mutant isolates throughout bioreactor cultivation as an exemplar.

## Discussion

In this study, we performed a transcriptomic meta-analysis to generate a high-quality gene co-expression network, and used this to predict biological processes for 9,579 (∼65%) of all *A. niger* genes including 2,970 hypothetical genes (ρ ≥ |0.5|). The compendium of resources developed in this work consists of (i) gene-specific sub-networks with two stringent Spearman cut-offs that ensure high confidence in biologically meaningful interpretations; (ii) statistically enriched GO terms for each co-expression network, and (iii) a refined list of co-expression relationships which incorporate over 1,200 verified ORFs to aid predictions of gene biological process based on experimental evidence.

In order to demonstrate the utility of these resources, we firstly interrogated these datasets at a gene-level, demonstrating that transcription factors VelC and StuA, which are critical components of ascomycete development and secondary metabolism, also regulate expression of the *A. niger* antifungal peptide AnAFP. Our data provide further evidence of the coupling between development, biosynthesis of secondary metabolites, and secreted antifungal peptides (24). The co-expression datasets can also be used to identify global regulators at the level of biological processes as demonstrated for fungal secondary metabolism and the two transcription factors TF1 and TF2 (which we name MjkA and MjkB). Generation of loss- and gain-of-function mutants demonstrated that they likely (in)directly regulate dozens of secondary metabolite loci at the transcript and metabolite level. Interestingly, MjkA is a Myb-like transcription factor highly conserved in Aspergilli with orthologues present in several plant genomes. Myb transcription factors have recently been demonstrated to regulate plant natural product biosynthesis (50), and our co-expression data and wet lab experiments suggest that titratable control of MjkA is a promising strategy for the activation of ascomycete secondary metabolism during drug discovery programs. With regards to the application of our co-expression approach to predict gene biological processes in other fungi, interrogation of the GEO database (23) demonstrates that several hundred global gene expression experiments are available for industrial cell factories (e.g. *A. oryzae, Trichoderma reesei*) and human or plant infecting fungi (e.g. *A. fumigatus, Cryptococcus neoformans, Candida albicans, Magnaporthe oryzae*), indicating that our approach can be broadly applied for industrial, medically, and agriculturally relevant fungi. As the financial costs for gene expression profiling continues to decline, this study paves the way for prediction of gene biological function using co-expression network analyses throughout the fungal kingdom.

## Availability of data and material

All datasets generated and/or analysed during the current study will be available in the FungiDB resource (http://fungidb.org/fungidb/).

## Acknowledgements

The authors wish to acknowledge the all members of the FungiDB project for providing the bioinformatics infrastructure to integrate the omic’s datasets of this study. Carina Feldle is acknowledged for her assistance during bioreactor cultivations.

## Authors’ contributions

P.S. performed the transcriptome meta-analyses, constructed the co-expression network and executed in silico quality analyses. M.J.K. analyzed sub-networks related to secondary metabolism. M.J.K., S.J. and N.P. generated deletion and overexpression strains and characterized them. M.J.K. and T.S. performed bioreactor cultivations. M.J.K. analyzed transcriptome and metabolome data derived from deletion and overexpression strains. B.B. and B.G. contributed to molecular analyses, B.N. and S.L. contributed to bioinformatics analyses. T.C. contributed to bioinformatics analyses and co-wrote the final text. V.M. initiated this study, coordinated the project and co-wrote the final text. All authors read and approved the final manuscript.

## References

1. Cairns TC, Nai C, Meyer V. How a fungus shapes biotechnology: 100 years of Aspergillus niger research. Fungal Biol Biotechnol [Internet]. 2018 May;5(1):13. Available from: https://doi.org/10.1186/s40694-018-0054-5

2. Show PL, Oladele KO, Siew QY, Aziz Zakry FA, Lan JC-W, Ling TC. Overview of citric acid production from Aspergillus niger. Front Life Sci [Internet]. 2015 Jul 3;8(3):271–83. Available from: https://doi.org/10.1080/21553769.2015.1033653

3. Pel HJ, Winde JH, Archer DB, Dyer PS, Hofmann G, Schaap PJ, et al. Genome sequencing and analysis of the versatile cell factory Aspergillus niger CBS 513.88. Nat Biotechnol [Internet]. 2007;25. Available from: http://dx.doi.org/10.1038/nbt1282

4. Gong W, Cheng Z, Zhang H, Liu L, Gao P, Wang L. Draft Genome Sequence of Aspergillus niger Strain An76. Genome Announc [Internet]. 2016;4(1):e01700–15. Available from: http://genomea.asm.org/lookup/doi/10.1128/genomeA.01700-15

5. de Vries RP, Riley R, Wiebenga A, Aguilar-Osorio G, Amillis S, Uchima CA, et al. Comparative genomics reveals high biological diversity and specific adaptations in the industrially and medically important fungal genus Aspergillus. Genome Biol. 2017;18(1).

6. Stajich JE, Harris T, Brunk BP, Brestelli J, Fischer S, Harb OS, et al. FungiDB: an integrated functional genomics database for fungi. Nucleic Acids Res [Internet]. 2011/11/09. 2012;40(Database issue):D675–81. Available from: http://www.ncbi.nlm.nih.gov/entrez/query.fcgi?cmd=Retrieve&db=PubMed&dopt=Citation &list_uids=22064857

7. Grigoriev I V, Nikitin R, Haridas S, Kuo A, Ohm R, Otillar R, et al. MycoCosm portal: gearing up for 1000 fungal genomes. Nucleic Acids Res [Internet]. 2014;42(Database issue):D699–704. Available from: http://www.ncbi.nlm.nih.gov/pubmed/24297253

8. Zerbino DR, Achuthan P, Akanni W, Amode MR, Barrell D, Bhai J, et al. Ensembl 2018. Nucleic Acids Res. 2018;46(D1):D754–61.

9. Meyer V, Andersen MR, Brakhage AA, Braus GH, Caddick MX, Cairns CT, et al. Current challenges of research on filamentous fungi in relation to human welfare and a sustainable bio-economy: a white paper. Fungal Biol Biotechnol [Internet]. 2016;3(1):1–17. Available from: http://dx.doi.org/10.1186/s40694-016-0024-8

10. Cerqueira GC, Arnaud MB, Inglis DO, Skrzypek MS, Binkley G, Simison M, et al. The Aspergillus Genome Database: Multispecies curation and incorporation of RNA-Seq data to improve structural gene annotations. Vol. 42, Nucleic Acids Research. 2014.

11. Cherry JM, Hong EL, Amundsen C, Balakrishnan R, Binkley G, Chan ET, et al. Saccharomyces Genome Database: The genomics resource of budding yeast. Nucleic Acids Res. 2012;40(D1).

12. Gov E, Arga KY. Differential co-expression analysis reveals a novel prognostic gene module in ovarian cancer. Sci Rep. 2017;7(1).

13. van Dam S, Võsa U, van der Graaf A, Franke L, de Magalhães JP. Gene co-expression analysis for functional classification and gene–disease predictions. Brief Bioinform [Internet]. 2017;bbw139. Available from: https://academic.oup.com/bib/article- lookup/doi/10.1093/bib/bbw139

14. Hsu C-L, Juan H-F, Huang H-C. Functional Analysis and Characterization of Differential Coexpression Networks. Sci Rep [Internet]. 2015;5(1):13295. Available from: http://www.nature.com/articles/srep13295

15. Dutkowski J, Kramer M, Surma MA, Balakrishnan R, Cherry JM, Krogan NJ, et al. A gene ontology inferred from molecular networks. Nat Biotechnol. 2013;31(1):38–45.

16. Cooper MA, Shlaes D. Fix the antibiotics pipeline. Nature [Internet]. 2011;472. Available from: http://dx.doi.org/10.1038/472032a

17. Macheleidt J, Mattern DJ, Fischer J, Netzker T, Weber J, Schroeckh V, et al. Regulation and Role of Fungal Secondary Metabolites. Annu Rev Genet [Internet]. 2016;50(1):371– Available from: https://doi.org/10.1146/annurev-genet-120215-035203

18. Boecker S, Grätz S, Kerwat D, Adam L, Schirmer D, Richter L, et al. Aspergillus niger is a superior expression host for the production of bioactive fungal cyclodepsipeptides. Fungal Biol Biotechnol [Internet]. 2018;5(1):4. Available from: https://fungalbiolbiotech.biomedcentral.com/articles/10.1186/s40694-018-0048-3

19. Meyer V, Ram AFJ, Punt PJ. Genetics, Genetic Manipulation, and Approaches to Strain Improvement of Filamentous Fungi. In: Manual of Industrial Microbiology and Biotechnology. 3rd Editio. NY: Wiley; 2010. p. 318–29.

20. Green MR, Sambrook J. Molecular cloningc: a laboratory manual. Vols. 1–3. Cold Spring Harbor, N.Yc: Cold Spring Harbor Laboratory Press; 2012. 1–2,028 p.

21. Meyer V, Wanka F, van Gent J, Arentshorst M, van den Hondel CA, Ram AF. Fungal gene expression on demand: an inducible, tunable, and metabolism-independent expression system for Aspergillus niger. Appl Env Microbiol [Internet]. 2011;77(9):2975– 19. 83. Available from: http://www.ncbi.nlm.nih.gov/pubmed/21378046

22. Fiedler MRM, Gensheimer T, Kubisch C, Meyer V. HisB as novel selection marker for gene targeting approaches in Aspergillus niger. BMC Microbiol [Internet]. 2017;17(1):57. Available from: http://bmcmicrobiol.biomedcentral.com/articles/10.1186/s12866-017-0960- 3

23. Barrett T, Wilhite SE, Ledoux P, Evangelista C, Kim IF, Tomashevsky M, et al. NCBI GEO: Archive for functional genomics data sets - Update. Nucleic Acids Res. 2013;41(D1).

24. Paege N, Jung S, Schäpe P, Müller-Hagen D, Ouedraogo JP, Heiderich C, et al. A transcriptome meta-Analysis proposes novel biological roles for the antifungal protein anafp in aspergillus Niger. PLoS One. 2016;11(11).

25. Gautier L, Cope L, Bolstad BM, Irizarry RA. affy—analysis of Affymetrix GeneChip data at the probe level. Bioinformatics [Internet]. 2004 Feb 12;20(3):307–15. Available from: http://dx.doi.org/10.1093/bioinformatics/btg405

26. Gentleman RC, Carey VJ, Bates DM, Bolstad B, Dettling M, Dudoit S, et al. Bioconductor: open software development for computational biology and bioinformatics. Genome Biol [Internet]. 2004 Sep 15;5(10):R80–R80. Available from: http://www.ncbi.nlm.nih.gov/pmc/articles/PMC545600/

27. Spearman C. The proof and measurement of association between two things. By C. Spearman, 1904. Am J Psychol. 1987;100(3–4):441–71.

28. Inglis DO, Binkley J, Skrzypek MS, Arnaud MB, Cerqueira GC, Shah P, et al. Comprehensive annotation of secondary metabolite biosynthetic genes and gene clusters of Aspergillus nidulans, A. fumigatus, A. niger and A. oryzae. BMC Microbiol [Internet]. 2013;13:91. Available from: http://www.pubmedcentral.nih.gov/articlerender.fcgi?artid=3689640&tool=pmcentrez&ren dertype=abstract

29. Sanchez JF, Wang CCC. The Chemical Identification and Analysis of Aspergillus Nidulans Secondary Metabolites. In: Fungal secondary metabolism - Methods and Protocols [Internet]. 2012. p. 97–109. Available from: http://link.springer.com/10.1007/978-1-62703-122-6

30. Park J, Hulsman M, Arentshorst M, Breeman M, Alazi E, Lagendijk EL, et al. Transcriptomic and molecular genetic analysis of the cell wall salvage response of Aspergillus niger to the absence of galactofuranose synthesis. Cell Microbiol. 26. 2016;18(9):1268–84.

31. Dobin A, Davis CA, Schlesinger F, Drenkow J, Zaleski C, Jha S, et al. STAR: Ultrafast universal RNA-seq aligner. Bioinformatics. 2013;29(1):15–21.

32. Love MI, Huber W, Anders S. Moderated estimation of fold change and dispersion for RNA-seq data with DESeq2. Genome Biol. 2014;15(12).

33. Benjamini Y, Hochberg Y. Controlling the false discovery rate: a practical and powerful approach to multiple testing. J R Stat Soc Ser B [Internet]. 1995;57(1):289–300. Available from: http://www.jstor.org/stable/2346101

34. Punt PJ, van Gemeren IA, Drint-Kuijvenhoven J, Hessing JG, van Muijlwijk-Harteveld GM, Beijersbergen A, et al. Analysis of the role of the gene bipA, encoding the major endoplasmic reticulum chaperone protein in the secretion of homologous and heterologous proteins in black Aspergilli. Appl Microbiol Biotechnol [Internet]. 1998;50(4):447–54. Available from: http://www.ncbi.nlm.nih.gov/pubmed/9830095

35. Hargrove TY, Wawrzak Z, Lamb DC, Guengerich FP, Lepesheva GI. Structure-functional characterization of cytochrome P450 Sterol 14alpha-Demethylase (CYP51B) from aspergillus fumigatus and molecular basis for the development of antifungal drugs. J Biol Chem. 2015;290(39):23916–34.

36. Scharf DH, Heinekamp T, Brakhage AA. Human and Plant Fungal Pathogens: The Role of Secondary Metabolites. PLOS Pathog [Internet]. 2014 Jan 30;10(1):e1003859. Available from: https://doi.org/10.1371/journal.ppat.1003859

37. Hoang HD, Maruyama JI, Kitamoto K. Modulating endoplasmic reticulum-Golgi cargo receptors for improving secretion of carrier-fused heterologous proteins in the filamentous fungus Aspergillus oryzae. Appl Environ Microbiol. 2015;81(2):533–43.

38. Dhingra S, Cramer RA. Regulation of sterol biosynthesis in the human fungal pathogen Aspergillus fumigatus: Opportunities for therapeutic development. Vol. 8, Frontiers in Microbiology. 2017.

39. Yasmin S, Alcazar-Fuoli L, Grundlinger M, Puempel T, Cairns T, Blatzer M, et al. Mevalonate governs interdependency of ergosterol and siderophore biosyntheses in the fungal pathogen Aspergillus fumigatus. Proc Natl Acad Sci U S A [Internet]. 2012;109(8):E497–504. Available from: http://www.ncbi.nlm.nih.gov/pubmed/22106303

40. Meyer V. A small protein that fights fungi: AFP as a new promising antifungal agent of biotechnological value. Vol. 78, Applied Microbiology and Biotechnology. 2008. p. 17–28.

41. Meyer V, Wedde M, Stahl U. Transcriptional regulation of the Antifungal Protein in Aspergillus giganteus. Mol Genet Genomics. 2001;266(5):747–57.

42. Hegedüs N, Sigl C, Zadra I, Pócsi I, Marx F. The paf gene product modulates asexual development in Penicillium chrysogenum. J Basic Microbiol. 2011;51(3):253–62.

43. Meyer V, Jung S. Antifungal Peptides of the AFP Family Revisited: Are These Cannibal Toxins? Vol. 6, Microorganisms. 2018.

44. Sheppard DC. The Aspergillus fumigatus StuA Protein Governs the Up-Regulation of a Discrete Transcriptional Program during the Acquisition of Developmental Competence. Mol Biol Cell [Internet]. 2005;16(12):5866–79. Available from: http://www.molbiolcell.org/cgi/doi/10.1091/mbc.E05-07-0617

45. Hu P, Wang Y, Zhou J, Pan Y, Liu G. AcstuA, which encodes an APSES transcription regulator, is involved in conidiation, cephalosporin biosynthesis and cell wall integrity of Acremonium chrysogenum. Fungal Genet Biol. 2015;83:26–40.

46. Clutterbuck AJ. A mutational analysis of conidial development in Aspergillus nidulans. Genetics. 1969;63(2):317–27.

47. Newman DJ, Cragg GM. Natural products as sources of new drugs over the 30 years from 1981 to 2010. Vol. 75, Journal of Natural Products. 2012. p. 311–35.

48. Nielsen KF, Mogensen JM, Johansen M, Larsen TO, Frisvad JC. Review of secondary metabolites and mycotoxins from the Aspergillus niger group. Vol. 395, Analytical and Bioanalytical Chemistry. 2009. p. 1225–42.

49. Bayram Ö, Krappmann S, Ni M, Jin WB, Helmstaedt K, Valerius O, et al. VelB/VeA/LaeA complex coordinates light signal with fungal development and secondary metabolism. Science (80-). 2008;320(5882):1504–6.

50. Liu J, Osbourn A, Ma P. MYB transcription factors as regulators of phenylpropanoid metabolism in plants. Vol. 8, Molecular Plant. 2015. p. 689–708.

